# PACE, Proximity-Associated Changes in Expression

**DOI:** 10.64898/2026.08.09.743800

**Authors:** Elijah Willie, Shreya Rajesh Rao, John Ormerod, Ellis Patrick

**Affiliations:** Centre for Cancer Research, The Westmead Institute for Medical Research, The University of Sydney; Sydney Precision Data Science Centre, The University of Sydney; School of Mathematics and Statistics, The University of Sydney; Charles Perkins Centre, The University of Sydney, New South Wales, Australia

## Abstract

Cellular transcriptional states are shaped by local tissue context, yet quantifying how cellular gene expression varies with proximity to different cell types remains challenging. Cell-resolved spatial transcriptomics data are typically sparse and susceptible to contamination from neighbouring cells through diffusion, imperfect segmentation and cell overlap, making it difficult to distinguish cell-state changes from technical artefacts. We present PACE (Proximity-Associated Changes in Expression), a hierarchical empirical Bayes framework for quantifying cell-type-resolved proximity effects on gene expression. PACE uses partial pooling to stabilise inference across genes and cell types, separates contamination from biological spatial associations, and identifies coordinated transcriptional programmes underlying each proximity effect. Applied to Xenium-profiled breast cancer tissue, PACE identifies stromal cells and macrophages as the populations whose expression varies most with tumour proximity, and resolves a shift in macrophage state at tumour interfaces. In a CosMx-profiled melanoma cohort, PACE characterises previously reported outcome-associated variation in macrophage SPP1 (osteopontin) by resolving its relationship with tumour proximity across clinical outcome groups. PACE provides an interpretable framework for quantifying how tissue organisation shapes cellular state in spatial molecular data.

## Introduction

Cells continuously respond to their tissue microenvironment through proximity-dependent mechanisms that underlie processes such as immune surveillance and tissue repair [Domínguez Conde et al., 2022, Liu et al., 2025, Arora et al., 2023]. Understanding how specific cell types influence the gene expression programmes of their neighbours is therefore central to interpreting tissue pathology and identifying therapeutic targets [Ma et al., 2023, Wang et al., 2023]. Spatial transcriptomic technologies such as Xenium [Janesick et al., 2023] and CosMx Spatial Molecular Imaging [He et al., 2022] now map RNA at single-cell resolution in intact tissue across diverse healthy and diseased contexts [Chen et al., 2020]. Realising this potential, however, requires computational frameworks that can quantify proximity-associated changes in gene expression while accounting for technical artefacts intrinsic to these platforms. Cell-resolved spatial transcriptomics data are inherently sparse, with only a small fraction of each cell’s transcriptome captured. In addition, transcript misassignment between neighbouring cells can arise through imperfect segmentation, RNA diffusion during tissue processing and *z*-axis projection artefacts [Saiselet et al., 2020, Pentimalli et al., 2025]. These contamination effects generate systematic false-positive associations between a cell’s expression and the identity of its neighbours that cannot be fully eliminated by improved segmentation alone [Ergen and Yosef, 2025]. Distinguishing proximity-associated transcriptional changes from these technical artefacts therefore remains a central analytical challenge in the analysis of spatial transcriptomic data.

Despite rapid methodological progress, existing approaches address only subsets of the challenges involved in quantifying proximity-associated changes in gene expression. Some methods quantify how local tissue context contributes to gene expression variation, for example by partitioning expression into intrinsic, environmental and interaction components [Arnol et al., 2019], relating expression to proximity to specific cell types [Iyengar et al., 2025], or modelling associations with neighbourhood composition [Mason et al., 2024]. Others move beyond individual genes to identify spatially patterned transcriptional programmes, allowing coordinated biological responses to be detected despite sparse transcript measurements [Tanevski et al., 2022, Dong et al., 2025]. A separate class of preprocessing approaches addresses transcript misassignment through computational decontamination and reassignment of incorrectly attributed molecules [Kwok et al., 2025, Marco Salas et al., 2025, Shi et al., 2026]. Nevertheless, no existing frame-work simultaneously resolves cell-type-specific proximity effects at the level of individual genes, identifies coordinated transcriptional programmes, accounts for transcript misassignment, and supports comparisons of proximity effects across samples and biological conditions.

Here we present Proximity-Associated Changes in Expression (PACE), a statistical frame-work for quantifying cell-type-resolved proximity effects on gene expression in cell-resolved spatial transcriptomic data. PACE stabilises inference across genes and cell types through hierarchical modelling, separates contamination from biological spatial effects, and identifies coordinated transcriptional programmes associated with each proximity response. In multi-sample studies, PACE separates shared proximity effects from condition-associated responses, supporting comparisons across biological and clinical settings. Together, these capabilities provide an interpretable framework for quantifying how local tissue organisation shapes cellular state. PACE is available as an R package from Bioconductor and as a Python package from PyPI.

## Results

To quantify how cells modulate their expression in response to specific neighbouring cell types, we developed PACE, a hierarchical empirical Bayes framework that decomposes gene expression variance into three interpretable components: baseline cell type identity, spatial cell state (proximity-associated expression changes), and technical contamination (Figure 1). Pairwise decomposition identifies which neighbouring cell types contribute most strongly to each focal cell type’s spatial cell state. Variance decomposition quantifies the relative contribution of baseline cell type identity, spatial cell state and technical contamination, while a per-gene driver score prioritises the genes underlying each focal–neighbour relationship.

**Figure 1:**
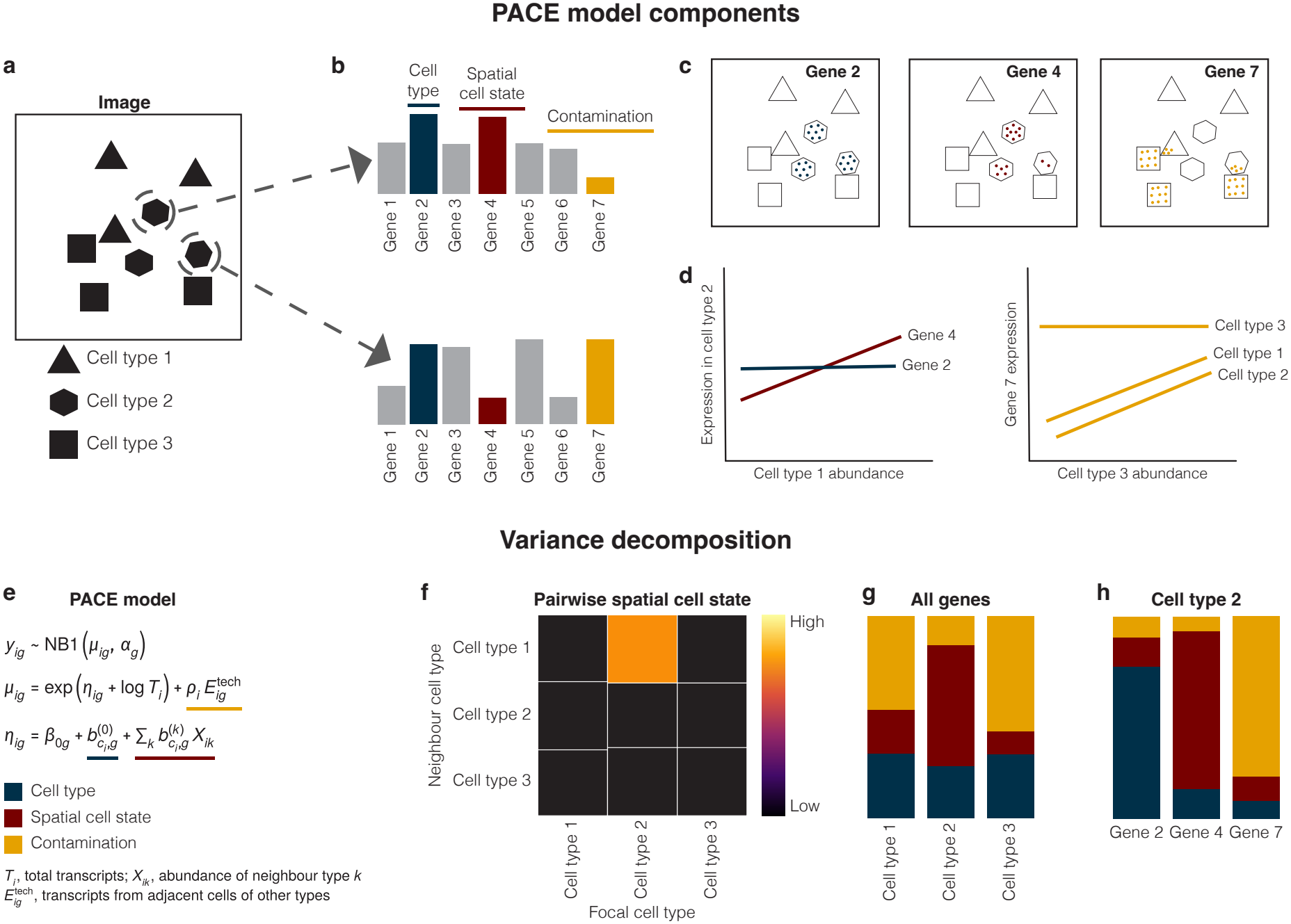
Overview of the PACE framework. (a) Schematic image with three cell types; two cells of type 2 are highlighted, one beside type 1 and one beside type 3. (b) Their expression profiles: gene 2 is high in both (cell type identity), gene 4 only beside type 1 (spatial cell state) and gene 7 only beside type 3 (contamination). (c) Transcript maps (dots) for genes 2, 4 and 7. Gene 4 transcripts are densest in type 2 cells closest to type 1, and gene 7 transcripts from type 3 cells are misassigned to the cells they touch. (d) Expression against neighbour abundance: gene 4 rises with cell type 1 abundance while gene 2 is unchanged, and gene 7 is constant in cell type 3 but rises in cell types 1 and 2. (e) The PACE model, with the cell type, spatial cell state and contamination terms underlined. (f) Pairwise spatial cell state heatmap of focal and neighbour cell types; here cell type 2 responds to cell type 1. (g, h) Variance decomposition by cell type across all genes (g) and by gene in cell type 2 (h), where genes 2, 4 and 7 are explained by cell type, spatial cell state and contamination respectively.

### PACE resolves tumour-associated spatial cell-state variation in breast cancer

We applied PACE to a human breast cancer Xenium section comprising approximately 126,000 cells spanning eight cell types to quantify the contribution of local tissue context to transcriptional variation.

The variance decomposition showed that, beyond the dominant cell type identity component, the spatial cell state component accounted for 0.3–5.8% of each focal type’s total expression variance and exceeded the technical contamination component (0.3–2.2%) in all populations except dendritic cells (Figure 2a). The spatial cell state component was largest in myoepithelial (5.8%), stromal (4.9%) and macrophage (4.2%) cells, indicating a measurable contribution of local cellular context to transcriptional heterogeneity in these populations.

**Figure 2:**
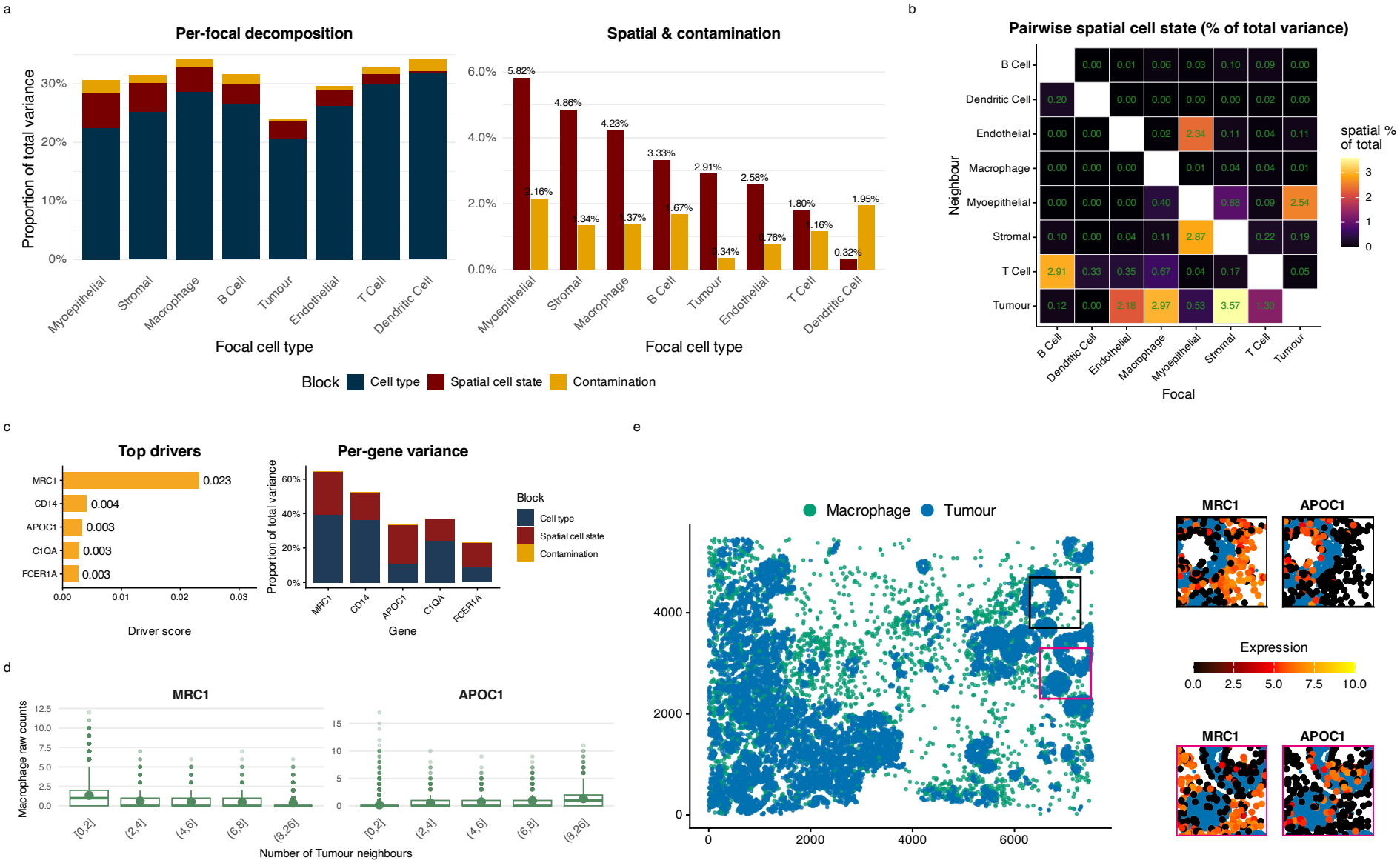
Tumour proximity is associated with a lipid-associated shift in macrophage state in breast cancer. (a) Per-focal variance decomposition across the eight displayed cell types. Left: full stacked bar showing the proportion of total expression variance attributable to cell type identity, spatial cell state, and contamination. Right: zoomed view of the spatial cell state and contamination components; the spatial cell state component is largest in myoepithelial, stromal and macrophage cells. (b) Pairwise spatial cell state heatmap: each entry gives the percentage of the focal type’s total variance contributed by proximity to that neighbour. Tumour as a neighbour contributes the most, with the stromal–tumour and macrophage–tumour pairs the two highest. (c) Macrophage–tumour drivers. Left: top genes ranked by driver score, with MRC1 the top-ranked driver. Right: per-gene variance decomposition for these genes, showing substantial spatial cell state contributions. (d) MRC1 and APOC1 expression in macrophages stratified by the number of tumour neighbours within 30 *µ*m. MRC1 decreases and APOC1 increases with tumour proximity. (e) Spatial map of the section showing macrophages (green) among tumour cells (blue), with all other cell types omitted for clarity. Paired insets show MRC1 and APOC1 expression in macrophages at two tumour-interface regions of interest (black and magenta boxes).

PACE next resolved the spatial cell state component of each cell type into specific focal– neighbour cell-type contributions. Tumour was the most influential neighbour type, with the stromal–tumour (3.6%) and macrophage–tumour (3.0%) pairings the two largest focal–neighbour contributions across the tissue (Figure 2b).

We next focused on the macrophage–tumour pairing to identify the genes underlying one of the strongest focal–neighbour spatial relationships. For each focal–neighbour pair, PACE can calculate a driver score which prioritises genes with large proximity-associated effects, high focal-cell specificity and sufficient expression in the focal population. The driver score ranked MRC1 (CD206) as the strongest spatial driver of the macrophage–tumour relationship, with lower expression in macrophages located near tumour cells (Figure 2c and 2d). Conversely, APOC1 increased with tumour-neighbour density, showing the opposite spatial pattern (Figure 2d). Visualising two tumour-margin regions of interest supported the spatial localisation of these gradients, with macrophages in tumour-dense regions showing reduced MRC1 and elevated APOC1, while those in tumour-sparse regions retained the opposite pattern (Figure 2e). MRC1 forms part of a tissue-resident macrophage programme, together with FOLR2 and LYVE1, associated with anti-tumour immunity, whereas APOC1 is co-expressed in lipid-associated macrophages linked to immunosuppressive tumour microenvironments [Nalio Ramos et al., 2022]. Together, these opposing gradients suggest that tumour proximity is associated with a shift in macrophage state, from tissue-resident features towards a lipid-associated programme. This spatial pattern is consistent with either phenotypic plasticity within individual macrophages or differential recruitment of macrophage subpopulations at the tumour boundary.

### PACE separates spatial signal from transcript contamination more effectively than existing methods

In dense tissues, transcripts from neighbouring cells can be misassigned to the focal cell through segmentation error, diffusion or cell overlap, so that spatial associations can reflect technical contamination instead of proximity-associated biology. PACE addresses this problem with a per-cell contamination term that absorbs the near-range ambient signal, allowing neighbour coefficients to more specifically capture biological proximity effects. To test whether this distinction affected gene prioritisation, we compared the top macrophage–tumour drivers from PACE with those from SpatioMark [Iyengar et al., 2025] and niche-DE [Mason et al., 2024]. Genes ranked highly by SpatioMark and niche-DE were predominantly canonical tumour genes, including ERBB2, KRT7, EPCAM and SCD, that were expressed far more strongly in neighbouring tumour cells than in the focal macrophages, consistent with contamination-driven prioritisation. By contrast, PACE prioritised macrophage-expressed genes, separating focal-cell spatial responses from neighbour-derived transcript signal (Figure 3a,b). This separation persisted after applying SpatioMark’s contamination correction, suggesting that removing an expression offset is insufficient when gene ranking does not account for focal-cell specificity. Across shared relationships, signed effects were concordant (Spearman *ρ* 0.74–0.94 in seven of eight pairs), so the methods differed mainly in which genes they prioritised, not in effect direction (Supplementary Figure S1).

**Figure 3:**
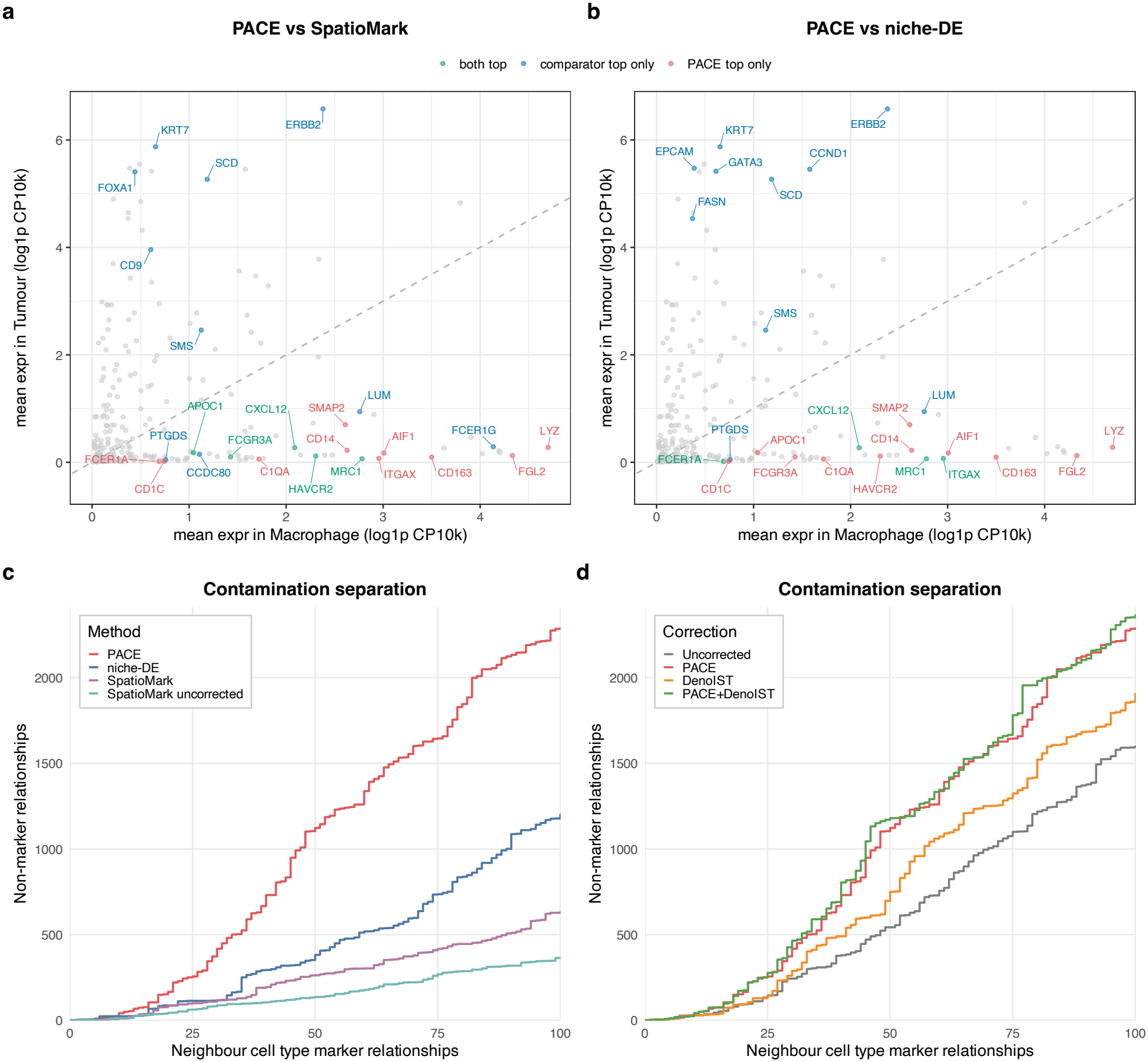
PACE separates proximity effects from transcript contamination more effectively than existing methods. (a–b) Macrophage–tumour driver ranking, PACE versus contamination-corrected SpatioMark (a) and PACE versus niche-DE (b). Each gene is plotted by its mean expression in macrophages (x) against neighbouring tumour cells (y) and coloured by which method ranks it in its top set. Genes above the diagonal are expressed more in tumour than in the macrophage, the signature of contamination. SpatioMark-top and niche-DE-top genes lie pre-dominantly above the diagonal, whereas PACE-top genes lie below. (c) Contamination separation across methods, each ranked on the relationships it returns. Ranking all relationships by significance, the curves accumulate non-marker (likely biological) against neighbour-marker (likely contamination) relationships. The methods compared are PACE, niche-DE, and SpatioMark, with SpatioMark shown both contamination-corrected and uncorrected. PACE recovers the most non-marker relationships per neighbour-marker relationship, and uncorrected SpatioMark the fewest. (d) The same separation across four correction strategies (uncorrected, PACE, DenoIST, PACE+DenoIST), plotted on the same axes as (c); PACE and PACE+DenoIST achieve the best separation.

We next asked whether this improved contamination separation extended across all inferred spatial relationships. Following the evaluation strategy used by SpatioMark [Iyengar et al., 2025], we ranked every gene–neighbour relationship by significance and labelled relationships in which the gene marked the neighbouring cell type as putative contamination false positives. Non-marker relationships were treated as putative biological positives, so better contamination separation corresponds to recovering more positives before accumulating neighbour-marker relationships. Each method was ranked on the gene–neighbour relationships it returned. PACE recovered 2,285 putative true-positive relationships at the first 100 false positives, compared with 1,178 for niche-DE, 628 for SpatioMark and 364 for uncorrected SpatioMark (Figure 3c).

Because count-level denoising also aims to reduce transcript misassignment, we tested whether PACE’s model-level correction provided additional benefit beyond upstream denoising. Using the same marker-based evaluation, we compared four correction strategies: no correction, PACE’s model-level contamination correction, count-level denoising with DenoIST [Kwok et al., 2025], and the combination of PACE and DenoIST. At the first 100 false positives, PACE recovered 2,285 putative true-positive relationships, a 43% increase over the uncorrected model (1,593, Figure 3d). DenoIST alone recovered 1,859 relationships (17% increase over uncorrected), whereas the combined PACE+DenoIST strategy recovered the most (2,352, 48% increase). PACE alone therefore outperformed DenoIST alone, and the two approaches appear complementary.

### PACE identifies condition-specific proximity effects in a multi-sample melanoma cohort

To demonstrate PACE in a multi-sample, condition-stratified setting, we applied it to a CosMx melanoma cohort of patients with progressive disease (PD) or non-progressive disease (non-PD, comprising stable and responding disease) following immune checkpoint blockade. In this setting, PACE included a responder spatial state component, which captures the part of each proximity effect that differs between clinical outcome groups. Variance decomposition showed that technical contamination was a substantial source of variation, comparable in magnitude to the cell type component in most focal populations and exceeding it in tumour cells (Figure 4a). Macrophages and fibroblasts carried the largest baseline spatial cell state contributions (0.85% and 0.62%, respectively), but these were much smaller than their corresponding contamination components (9.58% and 12.33%). The responder spatial state was smaller again, reaching at most 0.12% of total variance in fibroblasts. Outcome-dependent proximity effects were therefore small relative to both technical contamination and shared spatial cell-state variation.

**Figure 4:**
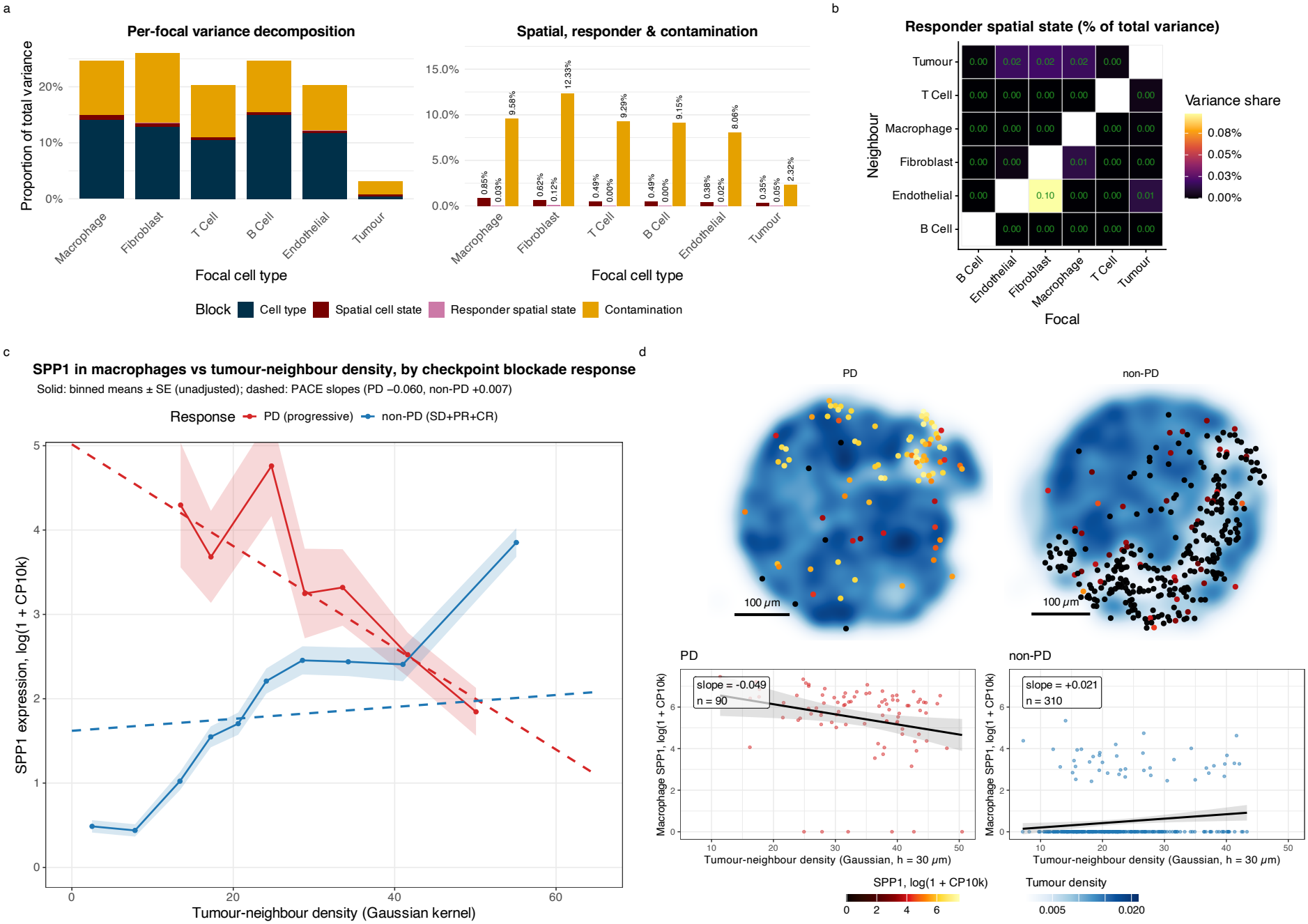
Macrophage SPP1 responses to tumour proximity diverge with immunotherapy out-come at the cohort level in melanoma. (a) Per-focal variance decomposition for the melanoma responder model. Left: full stacked bar. Right: zoomed view of the spatial cell state, responder spatial state, and contamination components. Macrophages and fibroblasts carry the largest spatial contributions; the responder spatial state is small across all populations, while contamination is highest in the sparsely distributed immune and stromal populations. (b) Pairwise responder spatial state heatmap: the percentage of the focal type’s total variance attributable to the (progressive disease *×* neighbour) interaction. (c) Macrophage SPP1 expression against tumour-neighbour density, split by immune checkpoint blockade response. Solid lines are binned means (*±*SE); dashed lines are the PACE spatial slopes (progressive disease*−*0.06, non-progressive +0.007), which diverge in sign. Binned means are unadjusted; PACE slopes are adjusted for contamination, image and the other neighbour types. (d) Exemplar images from a progressive-disease and a non-progressive-disease patient. Top: tumour-density map with macrophages coloured by SPP1 expression. Bottom: within-image SPP1 versus tumour-neighbour density, with the fitted slope and cell count; the progressive sample shows a negative slope (*−*0.049, *n* = 90) and the non-progressive sample a positive slope (+0.021, *n* = 310).

PACE next resolved the condition-dependent spatial component into specific focal–neighbour relationships and genes. Attributing this component to neighbour types (Figure 4b), responder-by-proximity effects were small throughout, the largest being in fibroblasts near endothelial cells (0.10% of total variance). For the macrophage–tumour interaction, SPP1 (osteopontin) was the only one of 927 genes with a significant responder spatial effect (slope *−*0.067, lfsr *<* 10^*−*4^; Supplementary Table S6). This agrees with the outcome-associated variation in macrophage SPP1 previously reported in this cohort with SIMVI [Dong et al., 2025] and extends it: PACE attributes the signal to proximity to tumour cells specifically, rather than to a learned neighbourhood representation, and estimates it after accounting for transcript contamination from adjacent tumour cells, which carry most SPP1 transcripts in this cohort. SPP1 marks an immunosuppressive tumour-associated macrophage population [Jiang et al., 2025, Lai et al., 2025]. Macrophage SPP1 showed different relationships with tumour proximity across outcome groups: in progressive disease, SPP1 expression declined with increasing tumour-neighbour density (slope = *−*0.06), whereas in non-progressive disease it was low away from tumour and the fitted slope rose only slightly with tumour proximity (slope = +0.007) (Figure 4c). Example images illustrated this cohort-level divergence, with SPP1-high macrophages located away from dense tumour regions in a progressive-disease sample (within-image slope = *−*0.049, *n* = 90) and near the tumour margin in a non-progressive sample (within-image slope = +0.021, *n* = 310) (Figure 4d).

### Runtime and sensitivity to the near-range radius

PACE is computationally efficient, with model fitting implemented in C++. Fitting the full breast cancer section (126,192 cells, 275 genes) took about 13 seconds on eight cores of a laptop (Apple M4, 24 GB memory), with wall-clock time scaling approximately linearly in the number of cells (Supplementary Figure S2c). Genes are processed in streaming chunks so peak memory scales linearly with cell number and independently of panel size. Runtime also grew approximately linearly with the number of genes: in the melanoma cohort (56,274 cells), a fit took about 2 seconds for 50 genes and 21 seconds for the full 927-gene panel. Additional cores gave a modest speedup at these panel sizes (about 2.2-to 2.3-fold from one to eight cores in both cohorts).

To assess sensitivity to the near-range radius used for contamination, we refitted both datasets across radii of 5–20 *µ*m. The primary proximity effects in breast cancer were essentially un-changed across this range, while contamination-marker relationships were progressively removed as the radius grew (Supplementary Figure S2a,b). In the melanoma cohort, the macrophage SPP1 responder divergence was stable across all tested radii (Supplementary Table S5), so the cohort-level estimate does not hinge on the exact radius choice.

## Discussion

We developed PACE, a hierarchical empirical Bayes framework for quantifying how cellular gene expression is shaped by the identity of nearby cell types, with explicit correction for transcript contamination inherent to cell-resolved spatial data. Applied to Xenium-profiled breast cancer and CosMx-profiled melanoma, PACE resolved proximity-associated transcriptional programmes consistent with known microenvironment remodelling, separating proximity effects from technical artefacts more reliably than SpatioMark and niche-DE and identifying condition-dependent spatial responses across a multi-sample cohort.

In the breast cancer dataset, macrophages near tumour showed reduced MRC1 alongside elevated APOC1 expression, consistent with the recently described axis of macrophage heterogeneity in breast cancer, where tissue-resident macrophages (MRC1, FOLR2, LYVE1) are functionally distinct from lipid-associated macrophages (APOC1, TREM2, APOE) [Nalio Ramos et al., 2022]. These tissue-resident macrophages, of which MRC1 is part, associate with CD8^+^ T cell infiltration and favourable prognosis, complicating the classical view of CD206 as a purely pro-tumour M2 marker [Modak et al., 2022, Ray et al., 2025].

In melanoma, PACE’s responder model resolved a condition-dependent divergence in macrophage SPP1 expression with tumour proximity between progressive and non-progressive disease. This is consistent with the role of SPP1 (osteopontin) as a marker of immunosuppressive tumour-associated macrophages, a population repeatedly associated with CD8^+^ T cell dysfunction, T cell exclusion, and poor immune checkpoint blockade outcome across solid tumours [Jiang et al., 2025, Lai et al., 2025]. In progressive disease SPP1 declines with tumour proximity, whereas in non-progressive disease it is maintained or rises. This cohort-level pattern agrees with the outcome-associated SPP1 variation previously reported with SIMVI [Dong et al., 2025], and PACE extends it by resolving the signal to the macrophage–tumour relationship, estimating how it differs between outcome groups, and separating it from contamination by adjacent tumour cells.

A broader motivation for this framework is the limited sensitivity of cell-resolved spatial transcriptomics. Even when genes are included in a targeted panel, transcript detection at the single-cell level is often sparse, making proximity-associated changes difficult to interpret from individual genes alone. PACE addresses this by estimating spatial effects across genes within a cell-type-resolved model, allowing weak and noisy gene-level signals to be summarised as interpretable variance components and then traced back to the focal–neighbour relationships and driver genes that explain them. This decomposition is useful because biologically coherent neighbourhood-associated programmes may be distributed across many imperfectly measured transcripts, not dominated by a few consistently detected genes.

The contamination correction operates at the model level rather than correcting individual transcript assignments, and can be combined with upstream count-level approaches such as DenoIST [Kwok et al., 2025] and DeLeakage [Shi et al., 2026]. In the breast cancer data, the combined PACE+DenoIST strategy performed slightly better than PACE alone, suggesting that model-level and count-level correction are complementary. However, the near-range term absorbs all signal within the specified radius, not only contamination, and contact-dependent mechanisms such as Notch–Delta interactions operating at very short range (*<*5 *µ*m) may be attenuated. The framework therefore prioritises specificity over sensitivity, which is appropriate when false-positive spatial associations are more problematic than missed short-range biology, but researchers investigating juxtacrine signalling should consider this trade-off.

The primary limitations of PACE each reflect the same underlying simplification. PACE models the tissue as a set of discrete, globally homogeneous, two-dimensional cell-type relationships. A fixed bandwidth cannot capture the full range of biological signalling scales, from direct juxtacrine contact to cytokine gradients extending well beyond 100 *µ*m, and the 2D projection may miss interactions between cells separated in the *z*-axis but within signalling range. Discrete cell type annotations smooth over continuous phenotypic variation, and a single global slope for each focal–neighbour pair assumes that an interaction operates with equal strength across tissue regions (e.g. the invasive margin and the tumour core). PACE also supports image-level random proximity slopes for between-image heterogeneity, but these need larger cohorts than those analysed in this study, so the condition-dependent effects reported here are cohort-level estimates. These simplifications are deliberate: they are what make the output interpretable. By committing to a structured model, PACE attributes variance, estimates effect sizes and identifies driver genes in ways that are harder to obtain from more flexible models. Relaxing these constraints while keeping gene-level interpretability is a natural next step.

## Methods

### Spatial neighbourhood quantification

PACE is a hierarchical empirical Bayes framework that decomposes cell-resolved spatial transcriptomics data into interpretable components: baseline cell type identity, proximity-associated transcriptional state, and technical contamination arising from transcript misassignment between adjacent cells. Cell-type-specific proximity effects are estimated as partially pooled random coefficients that share information across cell types through hierarchical regularisation, and contamination is absorbed by a per-cell coefficient estimated jointly with the biological model (Figure 1).

Separating biological proximity effects from contamination requires modelling two distance-dependent signals that operate at different spatial scales. Biological neighbourhood responses reflect cellular microenvironment composition at the scale of cell contacts and short-range paracrine signalling, whereas contamination arises from segmentation error, diffusion and cell overlap at much shorter range. PACE captures both using two distance-weighting kernels: a Gaussian biological kernel with bandwidth *h*_bio_ = 30 *µ*m that summarises the cellular neighbourhood, and a short-range exponential technical kernel with bandwidth *h*_tech_ = 5 *µ*m that captures the ambient contamination field.

For each focal cell *i* at spatial location **s**_*i*_ ∈ R^2^, the composition of its biological microenvironment is summarised by the kernel-weighted abundance of each cell type *k*,

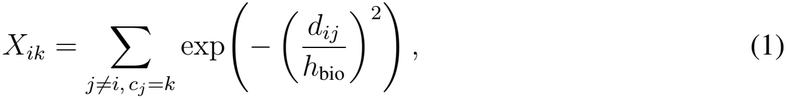

where *d*_*ij*_ = ∥**s**_*i*_ *−* **s**_*j*_∥ is the Euclidean distance between cells *i* and *j, c*_*j*_ ∈ *{*1, …, *K}* is the cell type of cell *j*, and *h*_bio_ = 30 *µ*m is the biological bandwidth. The sum is truncated at 3*h*_bio_ = 90 *µ*m and excludes the focal cell (*j≠ i*), giving an *n × K* matrix **X**. Neighbourhoods were constructed within each image for multi-sample cohorts and across the whole section otherwise.

### Per-cell contamination correction

Transcripts are misassigned between adjacent cells through imperfect segmentation, molecular diffusion, and projection of three-dimensional tissue onto the imaging plane [Marco Salas et al., 2025, Ergen and Yosef, 2025, Bai et al., 2021]. We modelled this as an additive per-cell contamination field. For cell *i*, the ambient field from its heterotypic neighbours is

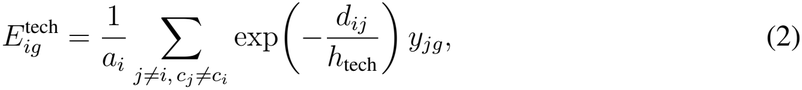

where *y*_*jg*_ is the count of gene *g* in neighbour *j, h*_tech_ = 5 *µ*m is the technical bandwidth, the sum is truncated at 3*h*_tech_ = 15 *µ*m, the default nuclear-expansion radius of Xenium cell segmentation [Marco Salas et al., 2025], and *a*_*i*_ is an isotropic area-fraction edge correction (the fraction of the disc within the imaged region). Homotypic neighbours are excluded (*c*_*j*_*≠ c*_*i*_). A per-cell coefficient *ρ*_*i*_, shared across genes, scales this field in the expected counts (equation 6) and is estimated jointly with the mean model.

The coefficient *ρ*_*i*_ is identified from anchor genes specific to other cell types, where this specificity is determined from expression in homotypic-core cells. For *N*_*i*_ = *{j* : *d*_*ij*_ *≤* 30 *µ*m*}*, the homotypic fraction

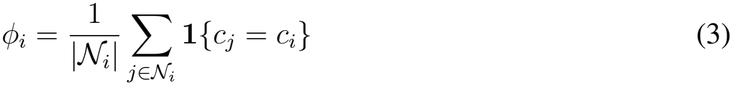

defines the core cells as those with *ϕ*_*i*_ *≥* 0.5. Let *m*_*cg*_ denote the mean of *y*_*ig*_ over the core cells of type *c*, with owner *o*(*g*) = arg max_*c*_ *m*_*cg*_. For a focal type *X*, the anchor set is

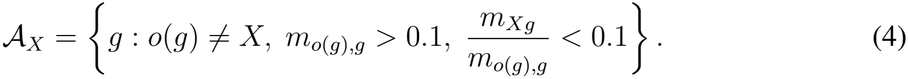

The strongest anchor genes for each focal cell type are listed in Supplementary Tables S3 (breast cancer) and S4 (melanoma).

### Hierarchical mixed models for gene expression

We modelled gene expression with hierarchical negative binomial regression, using a randomeffects specification across cell types as a regularisation device rather than a sampling assumption. Expression counts *y*_*ig*_ for cell *i* and gene *g* follow the NB1 (linear) parameterisation [Cameron and Trivedi, 1998], in which Var(*y*_*ig*_) = *µ*_*ig*_(1 + *α*_*g*_) for mean *µ*_*ig*_ and a gene-specific overdispersion parameter *α*_*g*_ estimated jointly with the mean model. The cell-type-specific coefficients are partial-pooling estimates that shrink towards zero, equivalent to empirical Bayes posterior modes under the Gaussian prior implied by the random-effects specification.

The count of gene *g* in cell *i* is modelled as

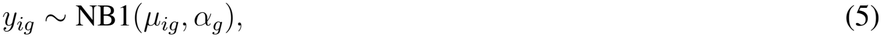

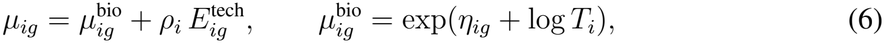

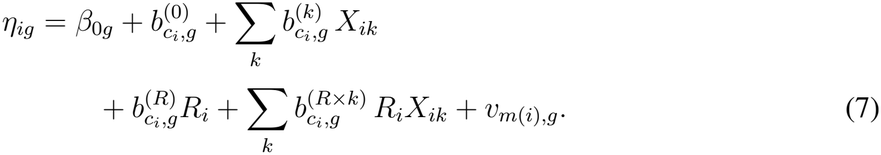

The expected count (equation 6) is an intrinsic biological mean 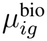plus the per-cell contamination 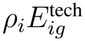. The biological mean carries a fixed per-cell exposure offset log *T*_*i*_, where *T*_*i*_ = ∑ _*g*_ *y*_*ig*_ is the total transcript count of cell *i*, so that proximity effects are estimated on expression rates, not raw counts; the additive contamination term is not scaled by *T*_*i*_. In the linear predictor (equation 7), *β*_0*g*_ is the global baseline of gene *g*, and the cell-type-specific coefficients, each given an independent Gaussian prior, are:

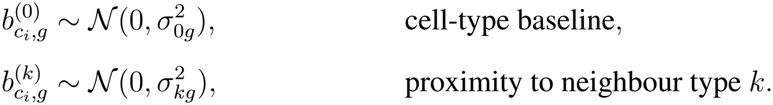

For multi-sample cohorts with a binary condition *R*_*i*_ ∈ *{*0, 1*}* (the condition indicator; here *R*_*i*_ = 1 for progressive disease following immune checkpoint blockade), two further terms enter the cell-type block,

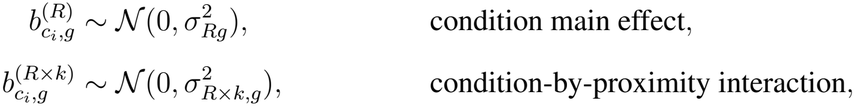

along with a per-image random intercept 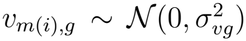, which absorbs image-to-image differences in baseline expression. Optionally, image-level random proximity slopes 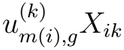, with 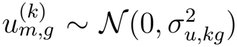, can be added to equation 7 to model between-image heterogeneity in proximity effects; these were not used in the analyses reported here. For a single tissue section the condition and image terms are omitted. All baseline and proximity coefficients are given independent Gaussian priors with cell-type-specific variance components, giving partial pooling across cell types.

### Model fitting

The additive contamination term 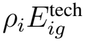 couples all genes through the shared loadings *ρ*_*i*_, and the cell-type-specific slopes act on continuous neighbourhood predictors across the full gene panel, so the model does not separate into independent per-gene fits. We therefore fitted it by penalised quasi-likelihood (PQL) [Breslow and Clayton, 1993, Breslow and Lin, 1995], cycling to convergence over the fixed and random effects, the per-cell contamination loadings *ρ*_*i*_, the gene-specific overdispersions *α*_*g*_, and the random-effect variance components.

The expected count 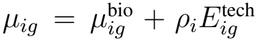, with biological mean 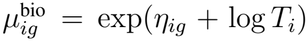, depends on the linear predictor only through 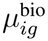, so the contamination term acts as a partial offset with 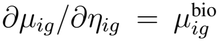. At each iteration a Gaussian working response and weight are formed for every cell and gene,

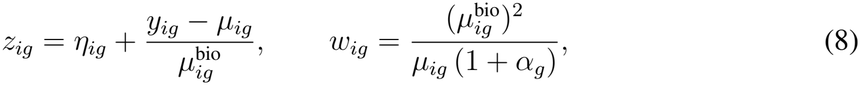

the NB1 weight following from Var(*y*_*ig*_) = *µ*_*ig*_(1 + *α*_*g*_). Treating *z*_*·g*_ as Gaussian with weights *w*_*·g*_, the fixed and random effects for gene *g* then solve the weighted mixed-model equations, with the cell-type coefficients and image intercepts penalised by their current variance components; this is a weighted least squares with a Gaussian prior on the random effects, returning their best linear unbiased predictors.

Given the current biological mean, the contamination loading *ρ*_*i*_ solves a weighted least-squares regression of the residual expression 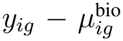 on the ambient field over the anchor genes 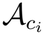 of cell *i*’s type, 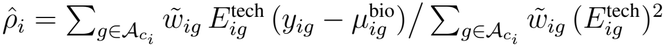. These anchor genes are expressed by the focal type at less than one tenth of the owning type’s level, so PACE approximates their intrinsic mean as zero 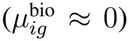. This is an approximation: contamination is itself a small fraction of the neighbouring cells’ expression, so an anchor’s residual intrinsic expression can be of the same order as its contamination signal. Under this approximation the estimator reduces to

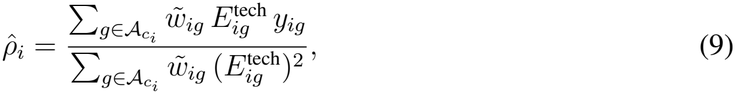

where 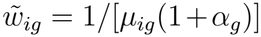is the negative-binomial precision, the inverse observation variance, distinct from the working weight *w*_*ig*_ of the mean-model update. The loading is constrained to be non-negative and empirical-Bayes shrunk towards a common value across cells, so that a cell with little anchor information borrows the population loading. Because anchor genes are expressed mainly by other cell types, the regression identifies *ρ*_*i*_ largely from technical signal, not from the biology it is meant to correct. Each overdispersion *α*_*g*_ is updated by maximum likelihood given the current mean, and the random-effect variances by an expectation-maximisation step. This step included a data-informed prior, enabled in all fits reported here. The prior scales each cell type’s proximity-slope variances by gene detection rate and neighbourhood-count variance, tightening these variances towards zero for sparsely supported focal–neighbour pairs. Through-out, genes are processed in chunks, so the full cell-by-gene working matrices are never formed; this reproduces the dense solution exactly, with peak memory scaling linearly in the number of cells and independently of the panel size.

### Empirical Bayes shrinkage and the local false sign rate

For each neighbour cell type, and each responder interaction in multi-sample cohorts, the proximity coefficients and their standard errors from the fit were stabilised jointly across focal cell types by multivariate adaptive shrinkage [Urbut et al., 2019, Stephens, 2017]. For a given neighbour, let 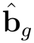collect the estimated proximity coefficients of gene *g* across the focal cell types, and **Ŝ** _*g*_ the diagonal matrix of their standard errors. The effects are modelled as draws from a mixture prior,

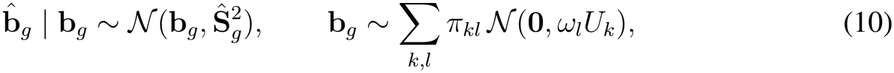

where the *U*_*k*_ are covariance components across the focal cell types, comprising canonical patterns together with data-driven components estimated from the strongest effects by extreme deconvolution, and *{ω*_*l*_*}* is a fixed grid of scaling factors. The mixture weights *π*_*kl*_ are estimate by maximising the marginal likelihood, so the procedure learns the panel-wide distribution of effects directly from the data while sharing information across focal cell types. Each posterior mean gives the shrunken estimates 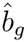(one per focal cell type) used in all downstream attribution and scoring, and their local false sign rate (lfsr), the posterior probability that the true effect is null or of opposite sign to the estimate [Stephens, 2017], quantifies confidence in the direction of effect. Genes with lfsr *<* 0.05 were taken to show a significant proximity effect.

### Variance decomposition framework

#### Per-gene decomposition

For each focal cell type and gene, we partitioned expression variance into cell-identity, spatial, contamination, and residual blocks in a single additive frame summing to one. Block magnitudes were computed from observed expression on the log(1 + CP10k) scale, with the within-cell-type partition fixed by the model’s fitted variance components [Nakagawa et al., 2017].

For focal cell type *c* and gene *g*, write *y*_*i*_ = log(1 + CP10k_*ig*_), with 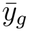the mean over all cells and 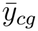the mean over cells of type *c*. The total variation of gene *g* within the focal population decomposes exactly into a between-cell-type and a within-cell-type term:

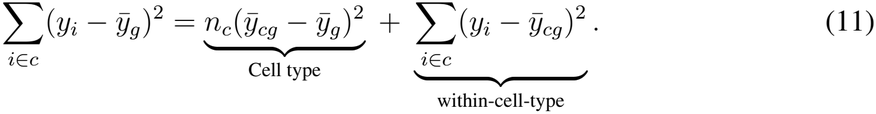

The Cell type block is the observed shift of the focal mean from the tissue mean. The within-cell-type sum of squares 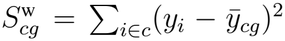 is apportioned to the Spatial, Contamination, and Residual blocks in proportion to the fitted per-gene variances of the proximity term (*V*_spatial_), contamination term (*V*_con_), and negative binomial dispersion (*V*_disp_),

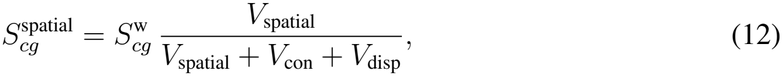

and likewise for Contamination and Residual. Here *V*_spatial_, *V*_con_ and *V*_disp_ are the variances, across the cells of type *c*, of the fitted proximity contribution 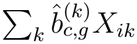, the fitted contamination contribution 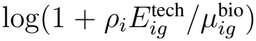, and the negative binomial dispersion, respectively. Because 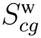 is the observed within-cell-type sum of squares, which already contains any covariance among these components, the proportional allocation distributes that covariance across the blocks without assuming the components are independent; the split is exact when the components are un-correlated and an approximation otherwise. For multi-sample cohorts the Spatial block is split into baseline and Responder spatial components in proportion to their fitted variances. The image random-effect variance and the responder main effect are retained in the fitted model but are not surfaced as separate decomposition blocks; their contribution to the within-cell-type sum of squares is apportioned across the reported blocks in proportion to the fitted variance components.

#### Multi-gene aggregation

Per-gene block sums-of-squares were pooled across genes following Goldstein, Browne and Rasbash [Goldstein et al., 2002],

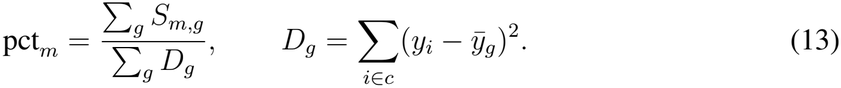

Mean-of-ratios and inverse-residual-variance-weighted alternatives identify the same three focal cell types with the largest spatial cell state component in breast cancer (Spearman *ρ ≥* 0.95 across focal types; not shown).

### Gene prioritisation

To rank the genes driving each cell type pair relationship, within each focal-neighbour pair (*f, h*) the driver score for gene *g* is

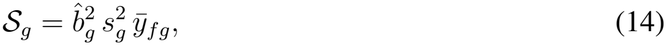

the product of its squared partially pooled proximity slope 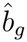for the pair, its squared specificity to the focal type 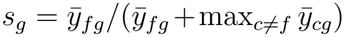, and its mean expression 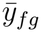in focal type *f*. Genes were retained at a local false sign rate below 0.05 and ranked within each pair.

### Pairwise attribution

The Spatial block of a focal cell type aggregates proximity effects from all neighbour types. To attribute it to individual neighbours we used the Pratt decomposition [Pratt, 1987], an additive allocation of a linear model’s explained variance to its predictors. The spatial component of the linear predictor for a focal cell of type *c* is 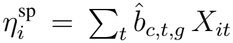, summing the partially pooled proximity slopes of gene *g* over neighbour types *t*. Writing Σ_*c*_ = Cov(**X**) over the cells of type *c* and 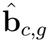for the vector of those slopes, the Pratt contribution of neighbour *t* for gene *g* is

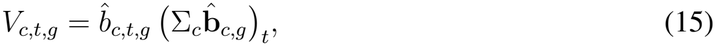

the product of the neighbour’s slope and its covariance with the fitted spatial predictor. Summing over neighbours recovers the spatial variance of gene *g* exactly, 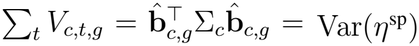, so the attribution is additive and leaves no residual. Pooling over genes, *V*_*c,t*_ = ∑ _*g*_ *V*_*c,t,g*_, gives the within-focal share 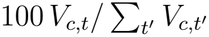; multiplying by the focal type’s Spatial block proportion yields the percentage of the focal type’s total variance attributable to neighbour *t*. The contributions are signed, and the small number of negative entries, where a slope opposes its co-variance with the spatial predictor, are set to zero for display. For multi-sample cohorts the same attribution was applied to the Responder spatial block.

### Contamination correction evaluation

To evaluate the per-cell contamination correction, we counted how many non-marker relationships each model recovers before accumulating false positives, treating markers of other cell types as false-positive controls, as established for contamination correction in imaging spatial transcriptomics [Iyengar et al., 2025, Kwok et al., 2025, Ergen and Yosef, 2025]. Cell-type markers were identified empirically by computing, for each gene, the area under the receiver operating characteristic curve of a one-versus-rest classification of cell types from the expression matrix; genes above a threshold (0.70) were designated markers of their cell type. The ordering of the methods, and the gain of PACE over the uncorrected model and over DenoIST alone, were unchanged at thresholds of 0.60 and 0.80. A (gene, neighbour) relationship whose gene marks the neighbouring cell type was treated as a false positive (a neighbour-marker relationship, indicating likely contamination), and all others as true positives (non-marker relationships, likely biological). Ranking every relationship by significance, as the absolute value of its test statistic (for PACE, the pre-shrinkage proximity coefficient divided by its standard error; for SpatioMark, its *t*-statistic; for niche-DE, the normal quantile of its two-sided *p*-value), and accumulating true against false positives, we compared each model (with and without the contamination term, alone and combined with count-level denoising) and each competing method by the number of true positives recovered at the first 100 false positives. Each model and each method was ranked on the relationships it returns.

### Evaluation data

We evaluated PACE on two cohorts, whose cell counts and cell-type composition are summarised in Supplementary Tables S1 and S2.

### Xenium breast cancer

We applied PACE to a single human breast cancer section profiled with Xenium [Janesick et al., 2023], using the nucleus-based cell segmentation supplied with the dataset, comprising 126,432 segmented cells with spatial coordinates and a 313-gene panel. Cells were annotated with sc-Classify [Lin et al., 2020], whose 17 transcriptional sub-states we collapsed into the modelled cell types. We excluded the rare mast population and retained genes detected in at least 5% of cells of some cell type, giving 126,192 cells across eight cell types (B cell, dendritic cell, endothelial, macrophage, myoepithelial, stromal, T cell, and tumour) and 275 genes. Because the data are a single section, neighbourhoods were computed across the whole tissue. The biological and technical kernels used bandwidths *h*_bio_ = 30 *µ*m and *h*_tech_ = 5 *µ*m.

### CosMx melanoma

We analysed a CosMx melanoma dataset of patients profiled after immune checkpoint block-ade [Dong et al., 2025]. We retained the 26 imaged sections of the 25 patients with a recorded best response (one patient contributed two sections) and contrasted progressive disease (PD; 7 patients, 7 sections, 13,161 cells) against non-progressive disease (the pooled stable, complete, and partial responders; 18 patients, 19 sections, 43,113 cells), giving 56,274 cells and 927 genes after detection filtering. Cells were annotated into six cell types (tumour, endothelial, fibrob-last, macrophage, B cell, and T cell); transcriptionally defined tumour subtypes were pooled into a single tumour class. Pixel coordinates were converted to micrometres by multiplying by 0.12028. Neighbourhoods were computed within each imaged section, and a per-section random intercept absorbed section- and patient-level differences in baseline expression. The kernels used *h*_bio_ = 30 *µ*m and *h*_tech_ = 5 *µ*m.

### Within-image associations

For display, within-image associations (Figure 4d) were summarised by the least-squares slope of log(1 + CP10k) expression on tumour-neighbour abundance *X*_*i*,tumour_ within each section, tested with a two-sided *t*-test at *α* = 0.05; these per-section tests are exploratory and were not adjusted for multiple comparisons.

## Data availability

Publicly available data were used for all evaluations, and were downloaded as described in the originating manuscripts. The Xenium breast cancer data are the public 10x Genomics human breast cancer dataset (In Situ Replicate 1) described in Janesick et al. [2023], available from 10x Genomics; the segmented and annotated SpatialExperiment analysed here is deposited at Zenodo. The CosMx melanoma data are described in Dong et al. [2025] and are available at Zenodo.

## Code availability

*PACE* is available as an R package from Bioconductor (development version), with development versions on GitHub, and as a Python package, `pace-spatial`, from PyPI, with source on GitHub and documentation at ecool50.github.io/pace-python. The two interfaces are built on the same compiled core, so an analysis may be carried out in either language. Both are released under the GPL-3 licence. All code for the analyses in this manuscript can be found on GitHub.

## Acknowledgements

The authors thank all their colleagues, particularly those at the Sydney Precision Data Science Centre and the Charles Perkins Centre, for their support and intellectual engagement. This work was supported by Australian Research Council Discovery Project grants to EP (DP260102836) and JO (DP210100521), and by the University of Sydney Postgraduate Excellence Award and the Charles Perkins Centre Jennie Mackenzie Research Fund to EW. The funding sources had no role in the study design; in the collection, analysis and interpretation of data; in the writing of the manuscript; or in the decision to submit the manuscript for publication.

## Author contributions

EW and EP conceived and designed the study. EW led the development of the PACE method and its statistical framework, with methodological input from JO and EP. EW implemented the R and Python packages, curated the datasets, and performed all data analysis. EW and EP drafted the initial manuscript, with writing contributions from SR, and all authors read and reviewed the text. JO and EP supervised the work and provided statistical guidance. All authors approved the final version.

## Use of AI tools

Claude (Anthropic) was used to assist with copy-editing. The authors take full responsibility for the content.

## Supplementary Material

### Supplementary Table: Dataset summary

Tables S1 and S2 summarise the two evaluation cohorts and their cell-type composition.

**Table S1:**
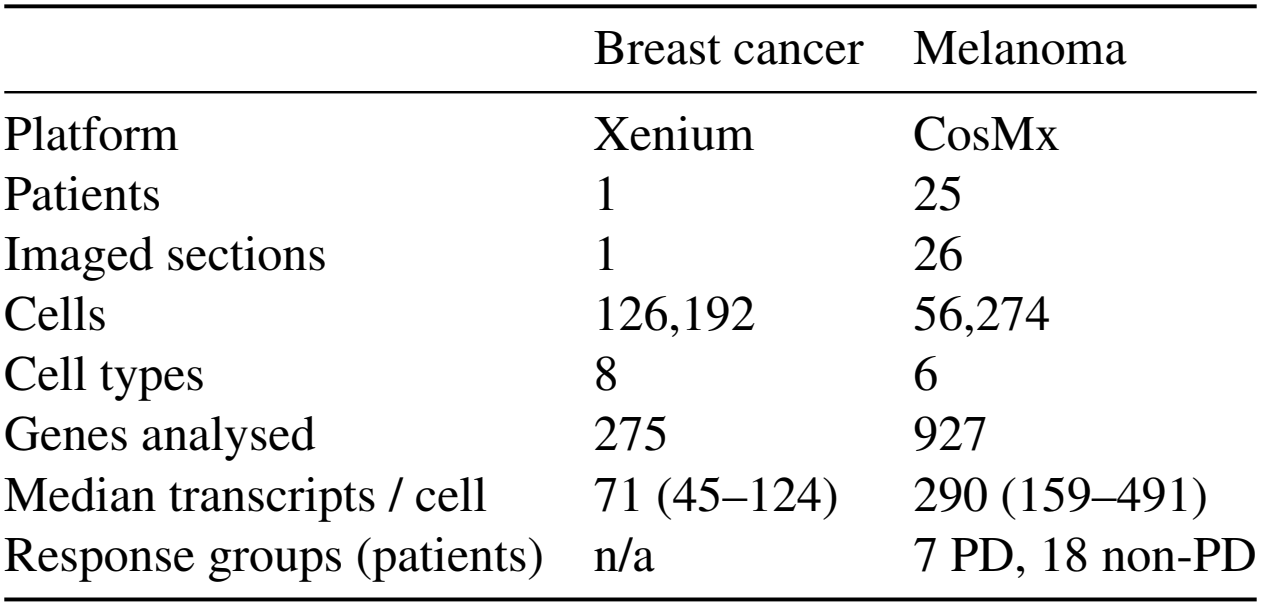
Summary of the two evaluation cohorts. Counts are the analysed cells that entered each fit; interquartile ranges are in parentheses.

|  | Breast cancer | Melanoma |
|---|---|---|
| Platform | Xenium | CosMx |
| Patients | 1 | 25 |
| Imaged sections | 1 | 26 |
| Cells | 126,192 | 56,274 |
| Cell types | 8 | 6 |
| Genes analysed | 275 | 927 |
| Median transcripts / cell | 71 (45–124) | 290 (159–491) |
| Response groups (patients) | n/a | 7 PD, 18 non-PD |

**Table S2:** Cell-type composition of each cohort: analysed cells and percentage of the cohort total.

| Breast cancer |  |  | Melanoma |  |  |
|---|---|---|---|---|---|
| Cell type | Cells | % | Cell type | Cells | % |
| Tumour | 59,606 | 47.2 | Tumour | 45,889 | 81.5 |
| T cell | 21,412 | 17.0 | T cell | 3,406 | 6.1 |
| Stromal | 13,250 | 10.5 | Macrophage | 3,101 | 5.5 |
| Macrophage | 11,243 | 8.9 | Fibroblast | 1,488 | 2.6 |
| Endothelial | 8,241 | 6.5 | Endothelial | 1,261 | 2.2 |
| Myoepithelial | 6,472 | 5.1 | B cell | 1,129 | 2.0 |
| B cell | 5,268 | 4.2 |  |  |  |
| Dendritic cell | 700 | 0.6 |  |  |  |

### Supplementary Table: Contamination anchor genes

Tables S3 and S4 list the strongest anchor genes used to identify the per-cell contamination loading in the breast cancer and melanoma cohorts respectively (Methods, “Per-cell contamination correction”).

**Table S3:** Top ten contamination anchor genes per cell type in the breast cancer cohort, ranked by expression in the owning cell type. Anchors are genes owned by another cell type and near-absent in the focal type, used to identify the per-cell contamination loading. The last column gives the total number of anchors for each type.

| **Cell type** | Top ten anchor genes | *n* |
|---|---|---|
| **B_Cell** | ERBB2, POSTN, LUM, ANKRD30A, KRT7, SCD, GATA3, CCND1, FOXA1, EPCAM | 164 |
| **Dendritic_Cell** | ERBB2, POSTN, ANKRD30A, KRT7, SCD, GATA3, CCND1, FOXA1, EPCAM, MLPH | 128 |
| **Endothelial** | ERBB2, ANKRD30A, KRT7, SCD, GATA3, FOXA1, EP-CAM, MLPH, TACSTD2, CXCR4 | 137 |
| **Macrophage** | ERBB2, ANKRD30A, KRT7, SCD, GATA3, CCND1, FOXA1, EPCAM, MLPH, TACSTD2 | 140 |
| **Myoepithelial** | POSTN, LUM, ANKRD30A, SCD, FOXA1, MLPH, CXCR4, FASN, LYZ, CXCL12 | 125 |
| **Stromal** | ERBB2, ANKRD30A, KRT7, SCD, GATA3, CCND1, FOXA1, EPCAM, MLPH, TACSTD2 | 145 |
| **T_Cell** | ERBB2, POSTN, ANKRD30A, KRT7, SCD, GATA3, CCND1, FOXA1, EPCAM, MLPH | 154 |
| **Tumour** | POSTN, LUM, CXCR4, MYLK, LYZ, AQP1, CXCL12, IL7R, PECAM1, SFRP1 | 148 |

**Table S4:**
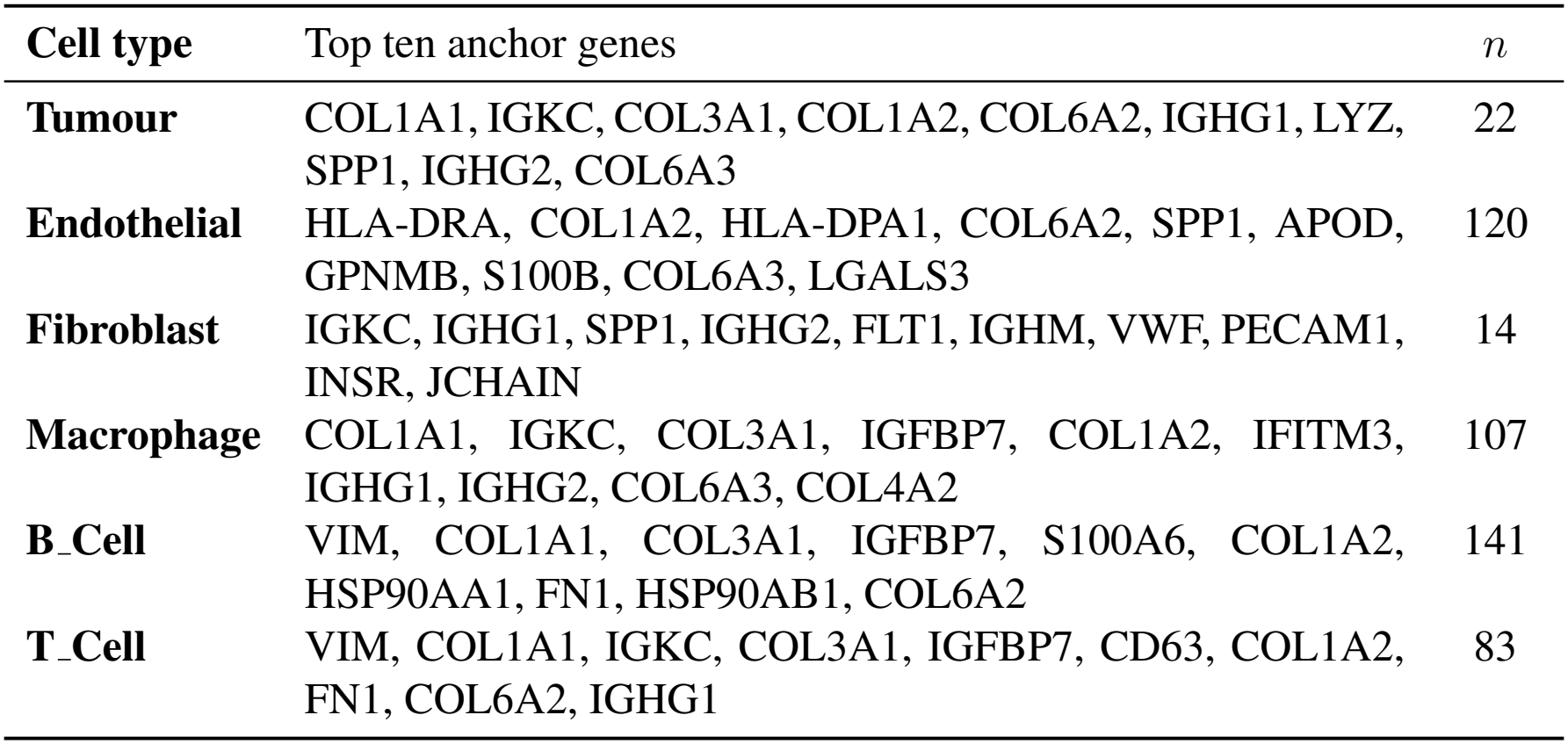
Top ten contamination anchor genes per cell type in the melanoma cohort, ranked by expression in the owning cell type. Anchors are genes owned by another cell type and near-absent in the focal type, used to identify the per-cell contamination loading. The last column gives the total number of anchors for each type.

| **Cell type** | Top ten anchor genes | *n* |
|---|---|---|
| **Tumour** | COL1A1, IGKC, COL3A1, COL1A2, COL6A2, IGHG1, LYZ, SPP1, IGHG2, COL6A3 | 22 |
| **Endothelial** | HLA-DRA, COL1A2, HLA-DPA1, COL6A2, SPP1, APOD, GPNMB, S100B, COL6A3, LGALS3 | 120 |
| **Fibroblast** | IGKC, IGHG1, SPP1, IGHG2, FLT1, IGHM, VWF, PECAM1, INSR, JCHAIN | 14 |
| **Macrophage** | COL1A1, IGKC, COL3A1, IGFBP7, COL1A2, IFITM3, IGHG1, IGHG2, COL6A3, COL4A2 | 107 |
| **B_Cell** | VIM, COL1A1, COL3A1, IGFBP7, S100A6, COL1A2, HSP90AA1, FN1, HSP90AB1, COL6A2 | 141 |
| **T_Cell** | VIM, COL1A1, IGKC, COL3A1, IGFBP7, CD63, COL1A2, FN1, COL6A2, IGHG1 | 83 |

### Supplementary Figure: Per-gene concordance with existing spatial methods

The main-text comparison (Figure 3) shows that PACE promotes fewer contamination-prone genes than niche-DE or SpatioMark. This reflects a difference in *ranking*, not a disagreement about the underlying spatial effects: where both PACE and a competing method call a (gene, neighbour) relationship significant, their signed effect estimates remain strongly concordant. Figure S1 quantifies this for the four tumour-as-neighbour pairs in the breast cancer section.

**Figure S1:**
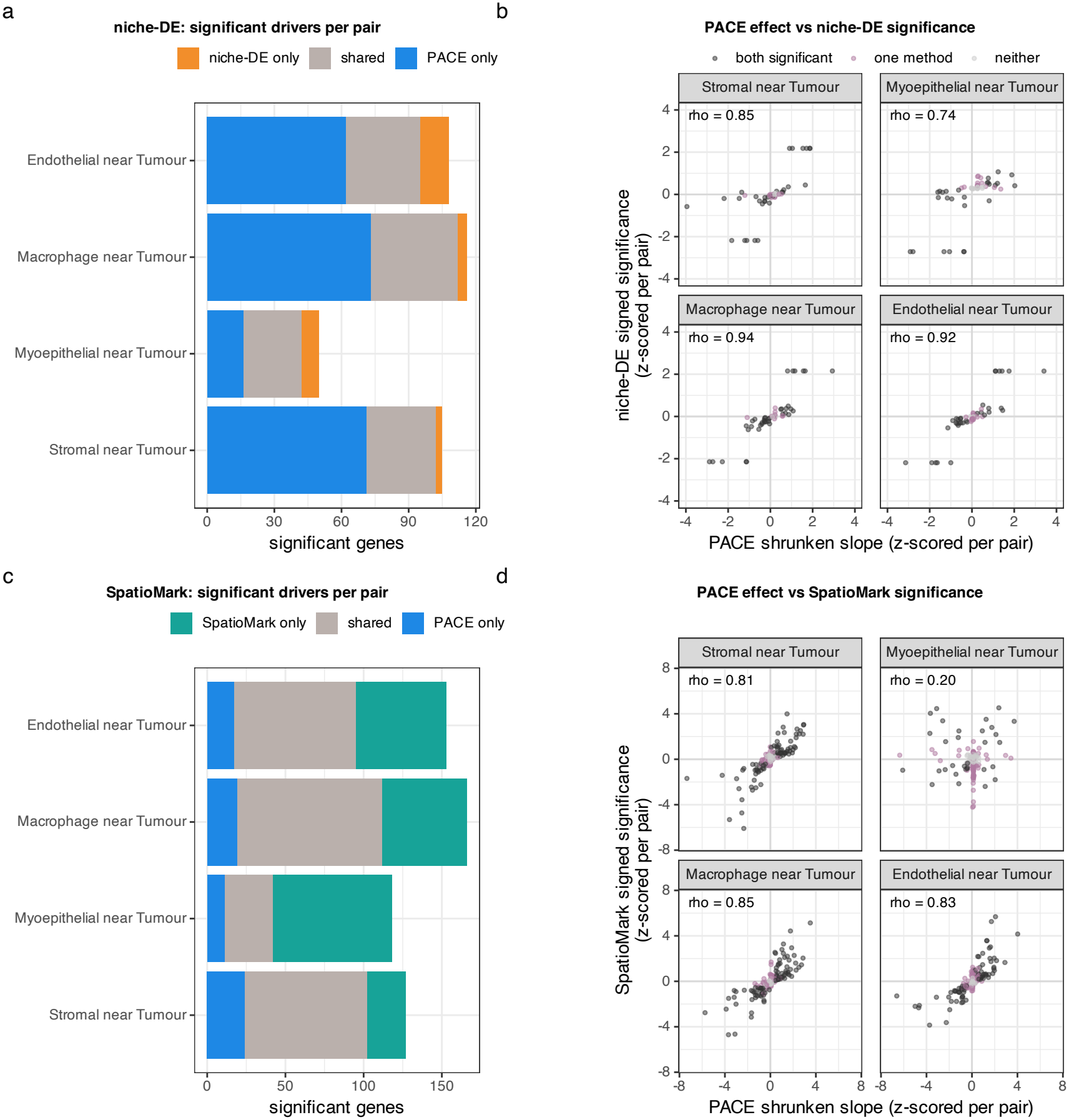
Per-gene concordance of PACE with niche-DE and SpatioMark in breast cancer. Top row, PACE versus niche-DE; bottom row, PACE versus the contamination-corrected SpatioMark arm. (a, c) Significant stromal/myoepithelial/macrophage/endothelial drivers of tumour proximity called by each method, split into method-only, shared, and PACE-only sets. niche-DE calls fewer genes than PACE (its specificity gate caps the tested set), whereas SpatioMark calls substantially more. (b, d) PACE shrunken slope versus the competing method’s signed significance, z-scored within each pair, coloured by whether both methods, one method, or neither call the gene significant. Per-pair Spearman correlations are high across nearly all pairs, indicating that the methods agree on the direction and rank of the spatial effects even though their top-ranked driver lists differ; the one exception, the myoepithelial–tumour pair under SpatioMark (*ρ* = 0.20), reflects contamination-driven relationships that PACE’s per-cell correction removes.

### Supplementary Figure: Near-range radius and runtime

The contamination kernel E^tech^ is truncated at three times its bandwidth, 15 *µ*m in the main analysis. Figure S2 shows how the macrophage–tumour results change when this radius is varied, and the runtime of a full fit.

**Figure S2:**
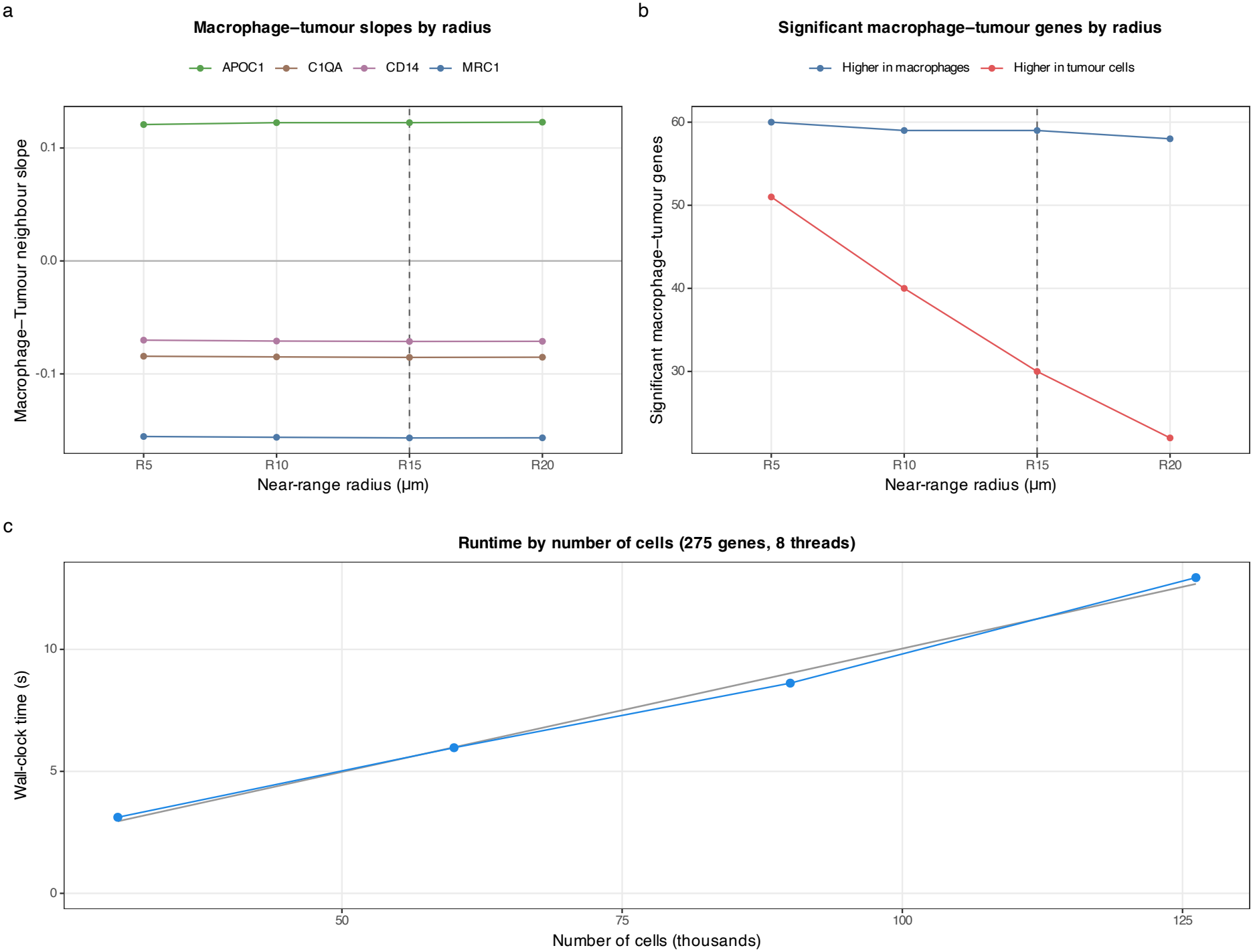
Near-range radius and runtime (breast cancer). (a) Macrophage–tumour slopes of MRC1, APOC1, CD14 and C1QA, refitted with near-range radii of 5–20 *µ*m. The slopes change little with the radius. (b) Number of macrophage–tumour genes significant at each radius (Bonferroni-corrected), split by whether the gene’s mean expression is higher in tumour cells or in macrophages. Genes higher in tumour cells are likely to reflect contamination; this split is simpler than the AUC-based marker definition used for Figure 3. As the radius increases from 5 to 20 *µ*m, the number of genes higher in tumour cells falls from 51 to 22, while the number higher in macrophages stays between 58 and 60. The dashed line marks the 15 *µ*m radius used in the main analysis. (c) Wall-clock time of a full fit against the number of cells (all 275 genes; eight cores of an Apple M4 laptop; 12 iterations). Runtime increases approximately linearly with the number of cells, and also with the number of genes (not shown): from 3.7 s for 50 genes to 12.4 s for 275 in breast cancer, and from 2.2 s for 50 genes to 21.0 s for all 927 in melanoma (56,274 cells).

### Supplementary Table: Melanoma near-range radius sensitivity

Table S5 refits the melanoma cohort across near-range radii of 5–20 *µ*m, confirming that the cohort-level macrophage SPP1 responder divergence is not an artefact of the 15 *µ*m setting used in the main analysis.

**Table S5:** Macrophage SPP1 slope with tumour-neighbour density in the melanoma cohort, refit across near-range (contamination-kernel) radii. The cohort-level outcome-dependent divergence in sign, positive in non-progressive and negative in progressive disease, is preserved at every radius; the 15 *µ*m setting is used in the main analysis.

| Near-range radius (µm) | Non-PD slope | PD slope |
|---|---|---|
| 5 | +0.008 | −0.046 |
| 10 | +0.007 | −0.058 |
| 15 | +0.007 | −0.060 |
| 20 | +0.007 | −0.059 |

### Supplementary Table: Melanoma responder-component gene ranking

Table S6 ranks the genes of the macrophage*←*tumour responder spatial component by local false sign rate, showing that SPP1 is the sole significant gene (lfsr *<* 0.05) among the 927 tested and that mashr shrinks every other condition-modulation slope to approximately zero.

**Table S6:** Gene-level ranking of the macrophage *←*tumour responder spatial component (the progressive-disease*×* tumour-neighbour interaction) in the melanoma cohort, ordered by local false sign rate (lfsr). The raw random-slope estimate is the pre-shrinkage PACE coefficient; the shrunken slope is the mashr posterior mean. SPP1 is the only gene reaching significance (lfsr *<* 0.05) out of 927 tested; mashr shrinks every other gene’s condition-modulation slope to approximately zero. Top 15 genes shown.

| Gene | Raw slope | Shrunken slope | lfsr |
|---|---|---|---|
| SPP1 | −0.067 | −0.067 | < 10<sup>−4</sup> |
| PSAP | +0.011 | 0.000 | 0.9992 |
| HLA-DPB1 | +0.011 | 0.000 | 0.9994 |
| C1QA | −0.008 | 0.000 | 0.9997 |
| HSP90AA1 | −0.009 | 0.000 | 0.9997 |
| GPNMB | +0.008 | 0.000 | 0.9998 |
| AZU1 | +0.008 | 0.000 | 0.9998 |
| CD63 | −0.008 | 0.000 | 0.9998 |
| TYK2 | +0.008 | 0.000 | 0.9998 |
| GPX1 | −0.007 | 0.000 | 0.9998 |
| MALAT1 | −0.007 | 0.000 | 0.9998 |
| CD74 | −0.007 | 0.000 | 0.9998 |
| LYZ | −0.007 | 0.000 | 0.9998 |
| FCER1G | −0.005 | 0.000 | 0.9999 |
| HSP90B1 | −0.006 | 0.000 | 0.9999 |

